# To preen or not to preen: aggressive and association networks predict allopreening interactions in wild parrots

**DOI:** 10.1101/2025.06.24.661407

**Authors:** Julia Penndorf, L. Fontana, J.M. Martin, L. M. Aplin

## Abstract

Allogrooming is fundamental to social relationships in many group-living mammals. In primates, allogrooming has been the subject of decades of research, and has been shown to play an essential role in maintaining social affiliations, and often interchanged for social tolerance and support. Yet, while the equivalent behaviour of allopreening occurs in many avian species, most description has been limited to its role in pair formation and maintenance, with little known about non-pair contexts. Here we investigated the decision-making associated with allopreening in wild sulphur-crested cockatoos, conducting detailed observations of 182 allopreening events between 126 individuals while concurrently recording rank, aggressive interactions, and social networks. We found no influence of sex as predicted if allopreening was primarily about pair bonding, but instead a positive correlation between allopreening and aggression (given and received). Within interactions, individuals were influenced by social association strength and rank, with individuals preening for longer and with more turn-taking when interacting with stronger social associates, and individuals investing more in interactions initiated by higher-ranked individuals. Our results are highly suggestive that allopreening may serve a convergent function in wild parrots to primates, both in maintaining social bonds and in negotiating social tolerance.

## Introduction

Allogrooming is considered a fundamental component of sociality in many mammals, including in primates, social mongoose, bats, and social rodents (Kutsukake & Clutton-Brock, 2006; Carter & Leffer, 2015), where individuals can spend up to one-fifth of their time combing through the fur of group members (Dunbar, 1991). Allogrooming is assumed to have significant hygienic benefits, such as the removal of ectoparasites that one can’t remove oneself (Stopka & Graciasova, 2001; Zamma, 2002; Mooring et al., 2004). Yet, in social primates, the total amount of time spent grooming is often better predicted by group rather than body size (a proxy for parasite load), suggesting that allogrooming provides benefits beyond the removal of parasites (Dunbar, 1991). For example, research on primates and other mammals has suggested that grooming interactions reduce stress levels (Boccia et al., 1989: Feh & de Mazieres, 1993; Gust et al., 1993: Wittig et al., 2008), play an important role in creating and strengthening social bonds (Mitani, 2009; Carter et al., 2020), and may increase fitness (Silk et al., 2003; Cameron et al., 2009; Silk et al., 2009).

Time budgets are limited, and the time individuals engage in social interactions must be traded off against other activities such as foraging or resting (Dunbar, 1992; Dunbar et al., 2009). Therefore, individuals should be selective regarding whom to groom. In Seyfarth’s (1977) classic model of primate grooming, the author proposed that individuals should groom up the hierarchy and those close in rank, in exchange for agnostic support or tolerance (Schino, 2001; Tiddi et al., 2012; Wubs et al., 2018). Further studies extended this model with biological market theory (BMT) to argue that grooming is a commodity to be traded for a range of benefits (Noe & Hammerstein, 1994, 1995). For example, grooming can be exchanged for tolerance (Carne et al., 2011; Borgeaud & Bshary, 2015; Garcia, Lemieux, et al., 2021), but also grooming (leading to reciprocal interactions Carne et al., 2011; Tiddi et al., 2011; Schino and Alessandrini, 2018), infant-handling (Gumert, 2007; Fruteau et al., 2011; Yu et al., 2013), mating opportunities (Norscia et al., 2009; Yu et al., 2013) or food-sharing (Carter & Wilkinson, 2013; Jaeggi et al., 2013). Evidence for both Seyfarth’s model and BMT remains mixed, but has been reported from primates (Tiddi et al., 2012), bats (Carter & Wilkinson, 2013), and meerkats (Kutsukake & Clutton-Brock, 2006)

In birds, allopreening (the equivalent of allogrooming) has been observed in over 400 species (Kenny et al., 2017), yet in contrast to mammals, allopreening has almost exclusively been described in two breeding contexts. First allopreening is commonly performed from parent to dependent offspring. Second, between adults allopreening plays an important role in courtship in strengthening bonds between mated pairs (Kushlan, 2011), in promoting cooperation for offspring feeding (Kenny et al., 2017) or the re-establishment of familiarity in mates after periods of separation (Erickson, 1973; Kushlan, 2011). Previous research has therefore highlighted that allopreening is more likely to occur in co-operatively or colonial breeding species, or in socially monogamous species with stable mate-bonds (Lewis et al., 2007; Gill, 2012; Kenny et al., 2017).

Some studies have suggested a wider, multifunctional social role for allopreening in birds, similar to that observed in social mammals. For example, at the group-level, allopreening may function to reduce aggression between neighbouring pairs in the colonially breeding common guillemot (*Uria aalga*), (Lewis et al., 2007), with reduced aggression in turn increasing breeding success. And at the individual level, dominant birds in groups of cooperatively breeding woodhoopoe (*Phoeniculus purpureus*) are more likely to receive preenings than subordinates (Radford & Du Plessis, 2006), and barn owl (*Tyto alba*) nestlings preen siblings in exchange for food (Roulin et al., 2016). By far most evidence for this multi-faceted role of allopreening in birds has come from corvids. For example, in common ravens (*Corvus corax*), allopreening events predict agonistic support (Fraser & Bugnyar, 2012), and in common ravens and carrion crows (*Corvus corone*), preening serves as reconciliation after aggressions with valuable interaction partners (Fraser & Bugnyar, 2011; Ikkatai et al., 2016; Sima et al., 2016; Sima et al., 2018). Yet, these studies predominantly focused on mixed-sex dyads, thus not fully removing potential effects of breeding context on preening.

There are multiple similarities between the social systems of many corvids and parrot species, and between these groups and primates (Fraser & Bugnyar, 2010). For example, all groups have been shown to form differentiated social bonds that are stable over time (corvids: reviewed by Boucherie et al., 2019, parrots: Aplin et al., 2021; Penndorf et al., 2023). Yet, perhaps surprisingly, allopreening in parrots has almost exclusively been described as between potential mates or with dependent offspring (Brockway, 1964; Garnetzke-Stollmann & Franck, 1991: Hobson et al., 2014—but see O’Connell et al., 2024, O’Connell et al., 2025) and largely studied for its role in pair formation (Rogers & McCulloch, 1981; Garnetzke-Stollmann & Franck, 1991). This view has been recently challenged in a comparative study in captive parrots and corvids showing that: (i) preening rates were similar to those measured in chimpanzees, (ii) preening interactions occurred outside the pair, and (iii) preening interactions were predicted by physical proximity and reduced aggression (Morales Picard et al., 2020). Yet, these results may need to be interpreted with caution, as data were collected in captivity, and artificial provisioning is highly likely to change the rate and the pattern of affiliative interactions (Garcia, Farine, et al., 2021).

In a previous study, we observed social allopreening in wild groups of sulphur-crested (SC-) cockatoos (*Cacatua galerita*), whereby individuals preferentially preened kin while also preening other unrelated individuals outside of the pair (Penndorf et al., 2023). SC-cockatoos are a large parrot, commonly observed across northern and eastern Australia that forms large stable roosting communities, with a minority of adults breeding in tree hollows near the main roost site, and roosting groups foraging in fission-fusion flocks in a shared home range (Penndorf et al., 2023). In this species, individuals form stable monogamous pair bonds (*personal observation*), but will also hold long-term, differentiated, social bonds outside the pair with both kin and unrelated individuals (Aplin et al., 2021). Individuals also exhibit evidence of social cognition in aggressive interactions, making decisions to initiate—or escalate—aggressive interactions based on rank (known individuals) or weight (unknown individuals; Penndorf et al., 2025).

The aim of this study is to test two predictions of Seyfarth’s model. First, as asymmetry in grooming interaction has been suggested to be correlated with tolerance (Noe & Hammerstein, 1994, 1995; Schino, 2001; Tiddi et al., 2012; Wubs et al., 2018), we tested whether the number of preening interactions within a dyad was predicted by the number of aggressions received or given. In alignment with previous studies (Noe & Hammerstein, 1994, 1995; Carne et al., 2011; Borgeaud & Bshary, 2015; Garcia, Lemieux, et al., 2021; Schino, 2001; Tiddi et al., 2012; Wubs et al., 2018; Morales Picard et al., 2020), we predicted the number of preening interactions to be negatively correlated with the number of aggressions received and given. Second, we tested whether the investment in preening interactions was predicted by the relative dominance rank of the participating individuals. In alignment with Seyfarth’s model, we predict preening interactions to be biased towards higher ranked individuals, with lower ranking individuals investing more into each interaction—both in terms of behavioural components and time spent preening—compared to higher ranking individuals.

## Methods

### Study population

Our study population comprised SC-cockatoos at four neighbouring roosting communities in northern Sydney, during winter in 2018 (one roost: BG), 2019 (three roosts: BA, CG & NB) and 2022 (two roosts: CG & BG-Figure 1a). At each roost site, individuals were attracted to the ground using small amounts of sunflower seeds. Once individuals were habituated, i.e. could be lightly touched on their back without inciting an adverse response, they were individually marked using non-toxic fabric dye (MARABU Fashion dye); (Penndorf et al., 2023) that lasted for three to four months. In addition, 144 birds had been ringed and wing-tagged with plastic cattle ear-tags as part of the ongoing—and unrelated—Citizen Science project *Big City Birds* (for details about the methods and tags see Davis et al., 2017), and some were present in the study population, as individuals regularly engage in localised between-movements (Aplin et al., 2021; Fehlmann et al., 2024). No birds were caught for this study.

**Figure 1:**
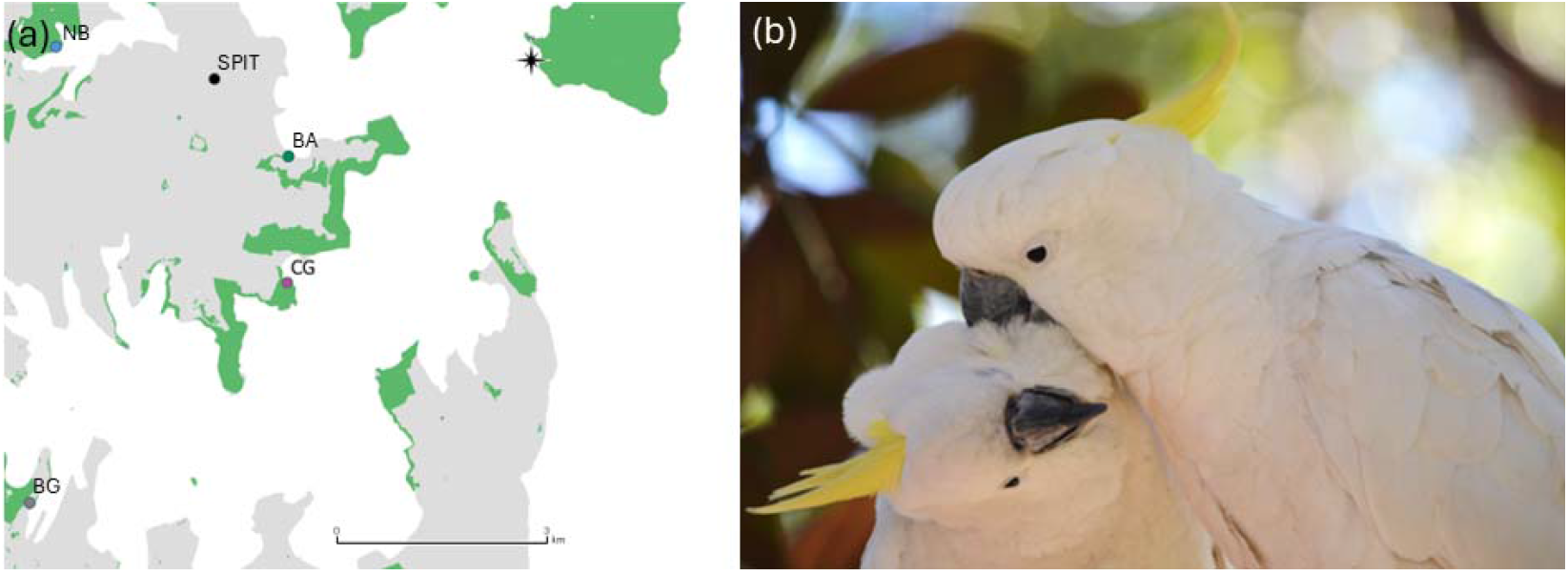
Study system and species. (a) Location of the roosting communities in Sydney included in this study. These include: Royal Botanic Garden (grey circle, BG—2018, 2022), Balmoral Beach (green circle, BA—2019), Clifton Gardens (violet circle, CG—2019, 2022) and Northbridge (blue circle, NB—2019). In addition, social associations were recorded at Spit Road (black circle, SPIT—2019), where individuals of all three northern roosts associated while being fed by a local resident Green areas represent parks and nature reserves, grey areas area build areas, and white is water. The map was constructed in QGIS (QGIS Development Team, 2009), using OpenStreetMap (OpenStreetMap contributors, 2017). (b) An example observation of two individuals allopreening.

Overall, between 136 and 536 uniquely recognizable birds were observed in each year of the study (2018: 136 paint-marked, 14 wing-tagged, 2 individually recognizable; 2019: 536 paint-marked, 17 wing-tagged, 8 individually recognizable; 2022: 233 paint-marked, 24 wing-tagged, 0 recognizable). Birds were aged by eye colour as juvenile (<7 years) or adult (>7 years), and amongst adults as male or female. In 2019 and 2022, feather samples were non-invasively collected, allowing verification of morphological sexing, and molecular sexing of juveniles. This also allowed re-identification of individuals between these two study periods.

### Social data collection

Social associations and interaction data were recorded in late winter, over 14 days in 2018 for a total of 42 hours in BG, over two 10 days periods for a total of 165 hours in 2019 (55 hours at BA, CG and NB respectively) and over 14 days for a total of 84 hours in 2022 (42 hours at BG & CG respectively; Figure 1a). Birds were attracted to begin ground foraging in parks close to the roost using small amounts of sunflower seeds scattered over a 300-680m^2^area (see Penndorf et al., 2023). The amount was variable depending on the size of the local night roost, ranging from 250g (NB) to maximum 450g (BA), depending on the size of the patch, and the number of birds present in the area. These food resources were non-monopolizable (sunflower seeds scattered over large areas), incentivizing birds to forage on the ground. Data collection only started when individuals foraged naturally (e.g., foraged on roots, grass, etc). Our previous study showed that networks collected in this context are highly clustered by roost membership (Penndorf et al., 2023).

Foraging flocks were then observed for 2.5 to 3 hours per day. During this time, *group scans* (Altmann, 1974) were conducted every ten minutes to record the identities of all birds present. Between scans, *all occurrence sampling* (Altmann, 1974) was used to record aggressive interactions and allopreening interactions. In all possible cases, the full sequence was recorded using pen and paper, including the initiator (hereafter *individual* 1), receiver (hereafter *individual* 2), and subsequent reciprocity (allopreening) or escalation (aggression). Additionally in 2022, a second observer (LF) filmed all preening interactions while other social interactions were recorded by JP. Allopreening interactions appear to be common at the communal roosts before dawn and after dusk. However, due to the difficulty of detailed observation in the roosting context (e.g., in tree canopies and low light conditions), we only focused only on allopreenings that were observed in the foraging context.

To make association networks, we constructed a *group-by-individual* matrix, where individuals were assigned to the same group if observed in the same presence scan. We then used the simple ratio index (SRI, R-package *asnipe*, Farine, 2013) to create an undirected association network for each year of the study. Edges between individuals were scaled from 0 (never associated) to 1 (always associated).

### Aggressive interactions

Overall, we recorded 9.908 aggressive interactions (BA_2019_=1.737, CG_2019_=2.735, CG_2022_=1.935, NB_2019_=1.005, BG_2018_=686, BG_2022_=1,810). These interactions were converted into a directed aggressive network for each year separately by the following formula:

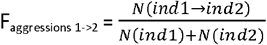

where Faggressions 1->2 is the frequency of aggression between individual 1 and individual 2, *Nind1->Ind2* the number of aggressions from individual 1 towards individual 2, *Nind1* the total number of aggressions initiated by individual 1, and *Nind2* the total number of aggressions initiated by individual 2. This result in a directed network, in which *Faggression* from 0 (individual 1 never aggressed individual 2) to 1.

In order to calculate a dominance hierarchy for each roosting community, we subset this data to include only individuals with more than ten aggressive interactions recorded at a given location (Sanchez-Tójar et al., 2018) and included all types of aggressive interactions in the analysis as simple ‘*wins*’ and ‘*losses*’. Dominance ranks were calculated using randomized Elo-ratings package aniDom version 0.1.5, (Farine & Sanchez-Tojar, 2021). Hierarchies were both linear and relatively stable over at least 3 years, with males tending to be dominant over females, and adults over juveniles, but with no other clear predictors of rank (Penndorf et al., 2024).

### Allopreening interactions

Altogether, we collated two types of allopreening data: (i) allopreening bouts including the initiator, receiver, and how often roles were exchanged, (ii) full durations of bouts and of episodes within bouts. Initiators were considered to be the individual first presenting to or preening the other individual (receiver).

### Analysis of social correlates of allopreening

To test initiation of preening interactions are correlated with social associations and/or aggressive interactions, we ran a Bayesian multi-membership model for proportions (zero-inflated beta binomial family, R-package brms Bürkner, 2017, Bürkner, 2018, 4 chains, 4,000 iterations). We modelled response variable—the preening rate of dyad *ij*—as binomial probability *p*_*ij*_ of individuals *i* and *j* being observed preening y times out of N_ij_ preening interactions initiated by individual *i*, allowing us to control for differences in the number of observed preening interactions individuals were involved in. The predictors were the social association index (SRI) and the proportion of aggressive interactions from individual 1 towards individual 2 (hereafter referred to as aggression given) and the proportion of aggressive interactions directed from individual 2 towards individual 1 (hereafter referred to as aggression received) during the entire study period.

### Analysis of investment in preening interactions

To investigate the investment of both individuals in an allopreening interaction, we considered: (i) the number of behavioural components (preenings) contributed by each individual to the interaction, and (ii) the total duration each individual preened its interaction partner during the interaction.

### Social predictors of the number of behavioural elements

For each individual, we quantified the number of behavioural elements contributed to the allopreening interaction. For example, if individual 1 preened individual 2, then individual 2 preened individual 1 before retreating, each individual would be considered having contributed one behavioural component to the interaction. To test whether the number of behavioural components contributed by each individual within the interaction was influenced by characteristics of the dyad, we then fitted multivariate Bayesian model (number of elements contributed by individual 1: Poisson family; number of elements contributed by individual 2: hurdle-Poisson family; R-package *brms*, Bürkner, 2017, 2018, 4 chains, 4,000 iterations, adapt_delta: 0.90), controlling for between-dyad variability (multi-membership model). Response variables were the number of behavioural components each individual contributed to a given interaction, with the initiator considered as individual 1 and the initial receiver involved in the preening was considered as individual 2. One of the predictors was the rank difference defined as the rank of the receiver minus the rank of the initiator, divided by the total number of individuals in the hierarchy. As previous work on parrots suggested that allopreening is more frequent between pairs (Brockway, 1964; Trillmich, 1976; Garnetzke-Stollmann & Franck, 1991; Hobson et al., 2014) and close-associates (Morales Picard et al., 2020), we additionally added the dyad was mixed or single sex (binary, TRUE/FALSE), and dyadic association strength (SRI). For the number of behaviours contributed by individual 1, we added an additional binary predictor of whether or not the receiver preened in return. The same predictor variables were used to predict the likelihood of individual 2 not reciprocating the allopreening.

### Social predictors of the allopreening duration

To test whether the investment of each individual-measured in time spend preening during the interaction-was influenced by characteristics of the dyad, we fitted a multivariate Bayesian model (shifted-normal family for preen_ID1_, and hurdle gamma family for preen_ID2_, R-package *brms*, Bürkner, 2017, 2018, 4 chains, 4,000 iterations, adapt_delta: 0.90). The response variable was the total preening duration given by the individual in seconds. Similar to the analysis for the investment in terms of behavioural components, the predictors were the rank difference, sex (mixed/same) and dyadic association strength (SRI). The same predictors were used in the hurdle part of the model, to predict the likelihood of non-reciprocation of the preening. For the duration individual 1 spent preening, we added an additional binary predictor, whether or not the receiver preened in return.

## Results

We recorded 182 allopreening events between 126 individuals (102 individuals present in one year only, 20 individuals sampled across 2 years each—Figure S4). Of the 182 allopreening events, 21% occurred in same-sex dyads, 34% in between-sex dyads, and 45% in dyads where the sex of at least one individual was unknown (Figure 2a). 42% of all preening interactions were directed up the hierarchy (Figure 1b), while only a minority of aggressive interactions were directed up the hierarchy (Figure 1c). We recorded the full preening sequence for 182 interactions: of those, most initiations involved an active allopreening of the receiver (92% of interactions), rather than a presentation (8% of interactions).

**Figure 2:**
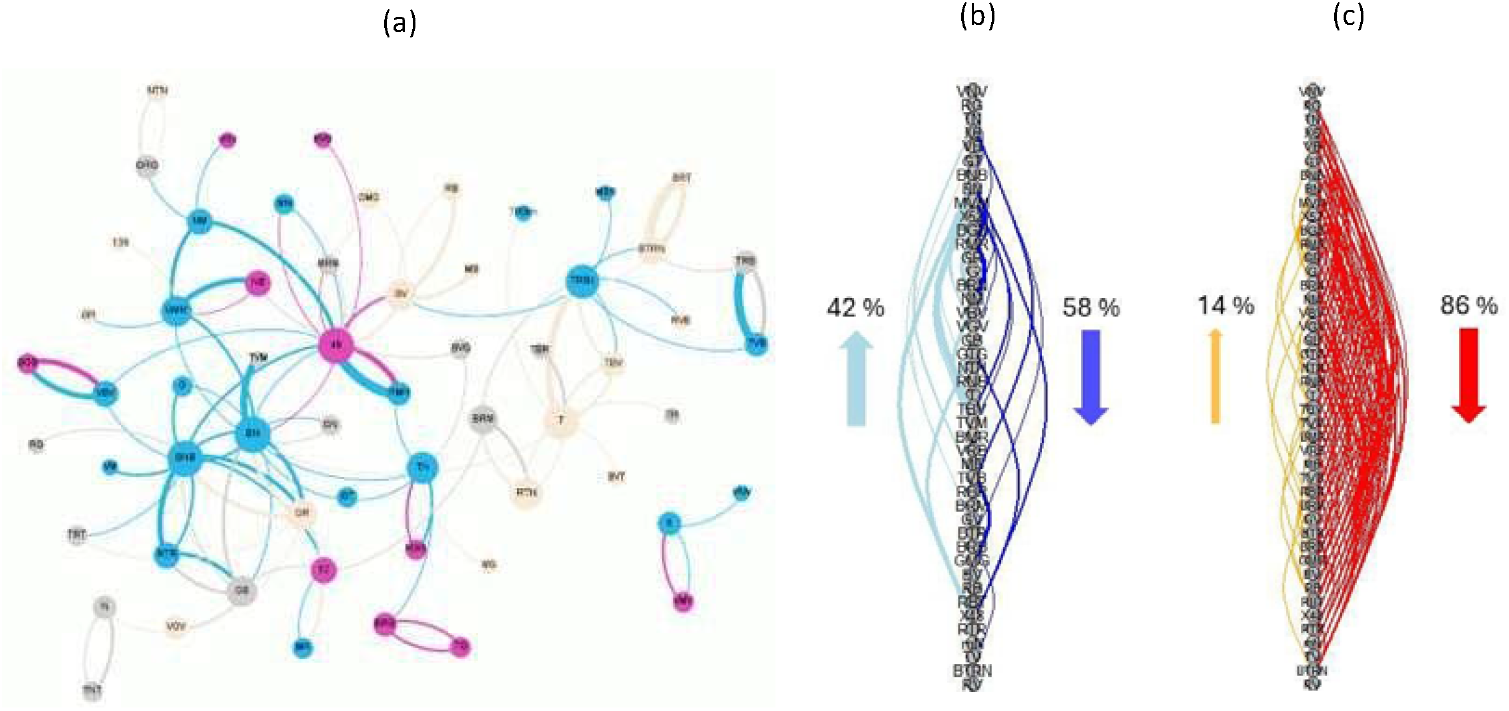
Interaction patterns in BG group (2018). (a) Directed allopreening network (n=85). Females are represented in pink, males in blue, juveniles in beige, and individuals of unknown age and sex in grey. Edges are coloured according to the direction of allopreening. (b) Aggressive interactions, and (c) allopreening interactions. Individuals are ordered according to dominance rank, from highest ranking (top) to lowest ranking (bottom). For each graph, darker edges on the right represent interactions directed down the hierarchy, while lighter edges on left are those directed up the hierarchy. Arrows surmise the relative proportion of interactions in each direction.

### Social predictors of allopreening

Perhaps counter-intuitively, individuals that exchanged more aggression were more likely to be observed allopreening. The initiation of allopreening was positively predicted by both aggressions directed towards (es= 18.61, se=6.59, 95% CI: [7.43,32.80]—Figure S1a), and received by (es= 26.20, se=9.22, 95% CI: [10.61,46.47]— Figure S1b), the receiver of the allopreening, even after accounting for large heterogeneity between dyads (es=2.95, se=0.67, 95% CI: [1.85, 4.47]). Association strength, however, was not correlated with the initiation of allopreening (es=0. 08, se=0.34, 95% CI: [-0. 61, 0. 75]).

### Social factors influencing the number of behavioural elements within a sequence

#### Number of elements contributed by the initiator

Within a single interaction, the length of the sequence was influenced by the association strength and the rank difference of the dyad, with initiators directing more behavioural elements towards weak associates (preen_ID1_: es=-0.78, se=0.38, 95% CI: [-1.50, −0.03]—Figures 3a & S3a), increasing their preening effort if the higher ranking individual was the first to preen, and with larger rank difference between the individuals (preen_ID1_: es= 0. 29, se=0. 15, 95% CI: [0, 0. 59]—97.4% of the posterior distribution was above zero; Figures 3a & S4a). Additionally, the number of behavioural elements contributed by the first individual was higher if it received preening(s) in return (preen_ID1_: es= 0.33, se=0.14 95% Cl: [0.05, 0.60]—Figure 3a). The sex of the dyad (same or mixed sex), however, did not influence the number of behavioural components contributed by the initiator (preen_ID1_: es= 0.07, se=0.15, 95% CI: [-0.23, 0.36]—Figure 3a).

**Figure 3:**
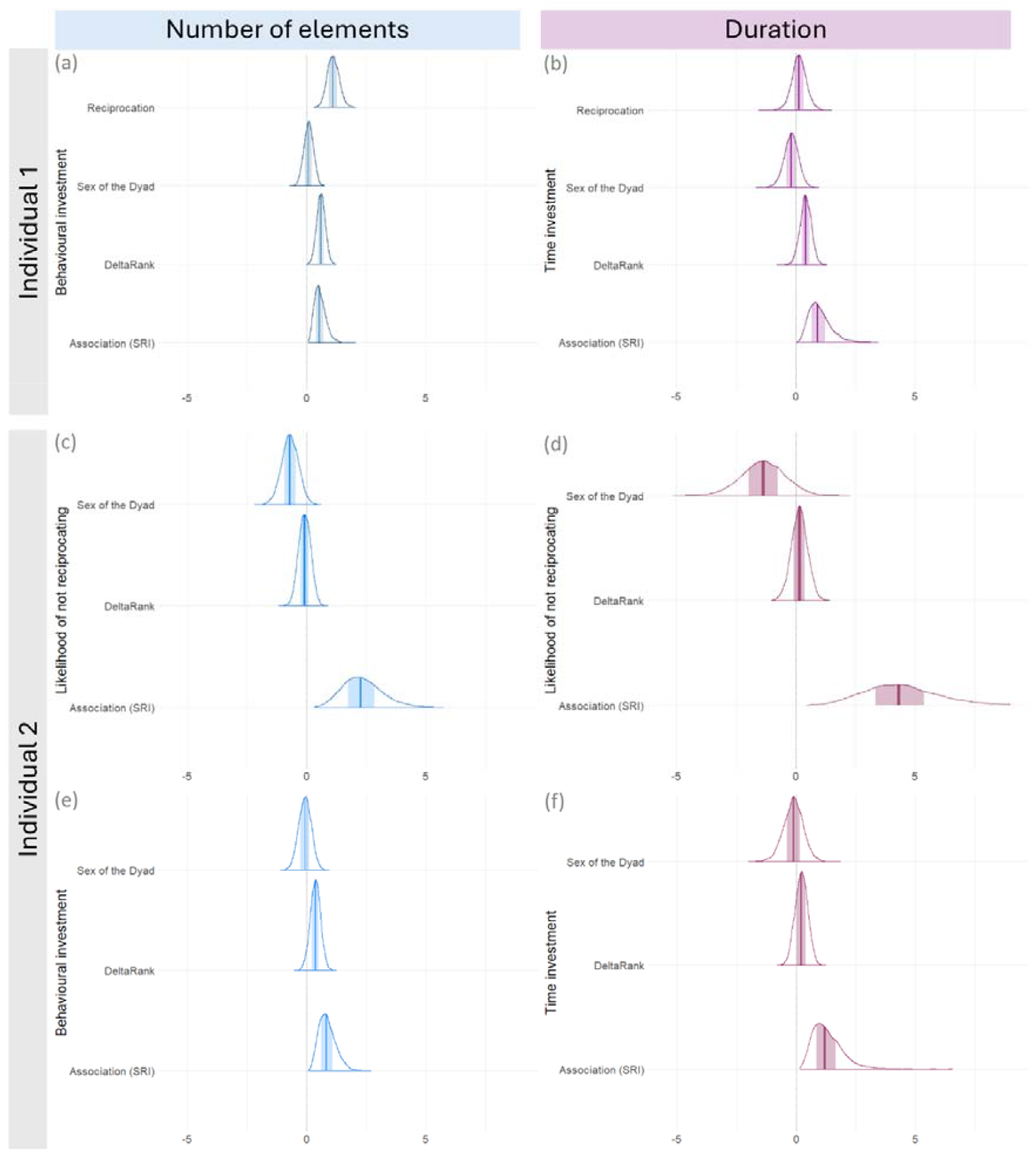
Posterior distributions with medians and 80% intervals. The left column (panels (a), (c) and (e)) correspond to the posterior distributions for analysis of the behavioural investments, measured as the number of elements each individual contributed to the preening sequence. The right column (panels (a), (c), (d) and (f)) represent the posterior distributions for the analysis of time investment, measured as duration each individual preened (in seconds). The first row (panels (a) and (b)) corresponds to the contributions of individual 1 (initiator). The second and third row (panels (c), (d), (e) and (f)) corresponds to the contributions of individual 1 (receiver). For each posterior distribution, the shaded area represents the 80% intervals, and the coloured line corresponds to the median.

#### Number of elements contributed by the receiver

Whether or not the receiver reciprocated the preening was best predicted by association strength and the sex of the dyad (same or mixed). Receivers were less likely to reciprocate preening(s) of close associates (preen_ID2_: es= 2.34, se=0.82, 95% Cl: [0.93, 4.10]—Figure 3c), and more likely to reciprocate interactions when in a mixed sex dyad (preen_ID2_: es= −0.70, se=0.35, 95% Cl: [-1.39,-0.04]—Figure 3c). The rank difference between the two individuals, however, did not predict the likelihood of reciprocation (preen_ID2_: es= −0.09, se=0.26, 95% CI: [-0.59, 0.41] —Figure 3c).

If the receiver reciprocated the interaction, the number of preenings was negatively correlated with the association strength (preen_ID2_: es= 0.86, se=0.35, 95% CI: [0.30, 1.66]—Figure 3e). Individuals tended to also increase their preening effort if the first individual to preen was higher ranked, and with larger rank differences between individuals (preen_ID2_: es= 0.36, se=0.21, 95% Cl: [-0.06, 0.77]—Figure 3e).

### Social factors influencing the duration of allopreening interactions

#### Duration of preening given by the initiator

Similarly to the analysis on behavioural components, the total duration of preening given by the initiator was negatively predicted by association strength (preen_ID1_: es= 0.98, se=0.44, 95% Cl: [0.32, 2.01]—Figures 3b & S3c) and rank difference, with the duration of preening given by individual 1 tending to increase with increasing rank difference (preen_ID1_: es= 0.41, se=0.24, 95% CI: [-0.07, 0.86]—Figures 3e & S4b). The sex of the dyad (mixed/same) did not influence the relative time investment of individual 1 (preen_ID1_: es= −0.19, se=0.30, 95% Cl: [-0.84, 0.37]—Figure 3b). Whether an interaction was reciprocated or not did not affect the duration given by the initiator (preen_ID1_: es=0.13, se=0.30, 95% Cl: [-0.46, 0.73]—Figure 3b).

#### Duration of preening given by the receiver

The reciprocation of a preening event was again best predicted by association strength, with individual 2 being less likely to reciprocate allopreenings of close associates (preen_ID2_: es=4.49, se=l.57, 95% CI: [l.84, 7.97]— Figure 3d). Additionally, the likelihood of individual 2 not reciprocating tended to be in lower mixed sex dyads (preen_ID2_: es=-1.39, se=0.96, 95% CI: [-3.29, 0.48]—Figure 3d). The rank difference between the dyad did, however, not affect the likelihood of individual 2 not reciprocating (preen_ID2_: es=0.15, se=0.96, 95% CI: [-0.52, 0.80]—Figure 3d).

If individual 2 reciprocated the preening, the total duration of the preening given by individual 2 was best predicted by the association strength of the dyad, with the duration of the preening increasing with decreasing association strength (preen_ID2_:es=l.29, se=0.62, 95% CI: [0.39, 2.77]—Figures 3f, S2d). Neither the sex of the dyad (same or mixed-preen_ID2_: es=-0.14, se=0.42, 95% CI: [-1.01, 0.67]—Figure 3f), nor the rank difference (preen_ID2_: es=0.21, se=0.26, 95% CI: [-0.31, 0.72]—Figure 3f), influenced the duration of the preening given by the receiver.

## Discussion

In previous work, we found that wild SC-cockatoos engaged in allopreening with multiple partners, with allopreening more likely to be observed between related individuals and individuals belonging to different roosts (Penndorf et al., 2023). Here we extend these results by using intensive observations over three years to examine the potential social functions of allopreening. We confirm our previous results and additionally find that: (i) birds with weak social associations exchanged longer preenings comprising more elements; (ii) dyads involved in more aggressive interactions also share more allopreening, (iii) contrary to Seyfarth’s model (which predicts increased preening effort between individuals close in rank) the investment of each individual—both in terms of number of elements contributed and overall duration—was positively correlated with rank difference, and (iv) in accordance with our prediction, the higher the rank difference, the more the preening sequence was biased towards the higher ranking individual.

Allopreening has been suggested to play an important role in maintaining social bonds (Spencer et al., 2005; Spoon et al., 2006; Kenny et al., 2017): individuals should therefore invest more into preening close associates to strengthen or maintain existing social bonds. This would result in either longer allopreening sequences or more frequent preening interactions between close associates. Intriguingly, our study found that individuals frequently observed together: (i) were not more likely to preen, and (ii) had fewer role exchanges within an interaction, while (iii) their allopreening interactions were of longer duration. This contrasts with previous studies in parrots, where preening frequency was positively correlated to association strength (Morales Picard et al., 2020). One explanation is that, in a foraging context, individuals preferentially direct interactions towards infrequent, but potentially valuable, interaction partners, such as for example kin (Penndorf et al., 2023) or high-ranking individuals. Alternatively, the more frequent, but shorter, exchanges between rare or infrequent associates could be a way for individuals to “test the waters” when forming new affiliative relationships. Under this scenario, role exchanges are expected to be more frequent (Carter et al., 2020). In our study, we do indeed find that the number of exchanged elements is highest for rare associates, likely representing a *quid pro quo* scenario, as the number of elements is higher if the exchange is reciprocated. Under this scenario, initial or rare interactions should also be relatively low-cost investments (O’Connell et al., 2025). While the observed pattern in our study is consistent with the “Testing the Waters” hypothesis (Carter et al., 2020, O’Connell et al., 2025), full confirmation would require further studies that track the full formation of relationships over longer periods of time.

Intriguingly, we did not find any evidence that preening sequences are longer between males and females as opposed to same-sex dyads. This could be an artefact of the protocol, as we recorded social interactions in a foraging context. Allopreening between pairs are frequent at their nest-hollow (*personal observation*), and social interactions are limited by feeding constraints (Garcia, Lemieux, et al., 2021). Therefore, within foraging associations, individuals may instead preferentially direct preening towards important social partners outside of the pair. Some evidence for this comes from a previous study on SC-cockatoos, suggesting that allopreening is maintained with kin even after dispersal (Penndorf et al., 2023). However, it is important to note that, by using sunflower seeds to attract birds to specific parks, we might have alleviated foraging constraints, and thus our dataset is likely to include an overrepresentation of rare interactions. Given that, for each site, less than 450 grams of sunflower seeds were scattered over a 300-680m^2^ area, and that birds spend most of their time foraging naturally (e.g., grass, roots), we believe this effect to be negligeable, however it remains to be tested.

Alternatively, individuals could favour allopreening with high-ranking individuals, as the latter may provide support or increase tolerance (Koyama et al., 2006; Carne et al., 2011; Tiddi et al., 2011; Borgeaud & Bshary, 2015; Garcia, Lemieux, et al., 2021). Our results also support this hypothesis in two ways. First, the investment of higher ranked individuals decreased with increasing rank difference. Second, the investment into the preening interaction increased if the interaction was initiated by the higher-ranking individual. Several studies have described rank bias within allopreening (allogrooming) interactions, with preening bouts being disproportionately directed towards the higher-ranking individual (Barrett et al., 2002; Schino et al., 2003; Kutsukake & Clutton-Brock, 2006; Radford & Du Plessis, 2006; Ventura et al., 2006; Miyazawa et al., 2020). Our study did not focus on the asymmetry of preening interactions, yet we provide intriguing insight on how rank differences influence affiliative interactions.

When considering the wider pattern of preening interactions, we found a positive correlation between preening and aggression. While perhaps initially surprising, this result is similar to that described in several primate species (Barrett et al., 2002; Schino et al., 2003; Port et al., 2009) and meerkats (Kutsukake & Clutton-Brock, 2006). One potential explanation proposed for the positive correlation between preening and aggression has been the need for reconciliation with valuable interaction partners. Reconciliation after aggressive interaction has been shown to occur in corvids (Fraser & Bugnyar, 2011; Sima et al., 2018), and has been suggested to occur in captive budgerigars *Melopsittacus undulatus* and monk parakeets *Myiopsitta monachus* (Morrison, 2009; Ikkatai et al., 2016). An alternative, but non-exclusive, explanation is that such patterns may emerge when subordinates invest in preening (grooming) to appease potential aggressors (Schino et al., 2003), exchanging preening (grooming) for future tolerance. This second explanation is consistent both our findings of unbalanced reciprocity in preening and of a correlation between preening and aggression networks. If so, this is suggested to primiarly occur only within dyads close-by in rank (Roubova et al., 2015). In this case, our limited sample size restricted a detailed analysis of this question. However, our recent work has identified that SC-cockatoos use strategic heuristics when choosing where to direct their aggressions (Penndorf et al., 2025). So, it seems reasonable that allopreening may also be used to consolidate or improve rank, or in exchange for tolerance, in addition to its affiliative role strengthening bonds within mates and close kin (Penndorf et al., 2023). Fully disentangling these two hypotheses (allopreening as reconciliation vs allopreening for tolerance) would require a time-sequence analysis between allopreening (grooming) and aggressive interactions, as well as experiment testings whether allopreening (grooming) improves the subsequent tolerance (such as, for example, done by Garcia, Lemieux et al., 2020). While this was beyond the scope of our study, it constitutes an exciting future direction.

If allopreening is exchanged for tolerance, this might further suggest the intriguing possibility of the presence of a biological market in SC-cockatoos. This, to our knowledge, would be the first such description in a wild parrot. Yet, preening being directed up the hierarchy is, by itself, not support for a biological market. Further evidence is needed that commodities are exchanged, and that the outcome of the exchange is affected by the availability of traders on the market (Port et al., 2009). For example, tolerance may serve as commodity in Vervet monkeys as dominant individuals increase their tolerance (measured as in the distance at which individuals foraged on artificially provided food) after being groomed by subordinates (Borgeaud & Bshary, 2015; Garcia, Lemieux, et al., 2021). While evidence is currently lacking in this case, support for the biological market theory could be provided by identifying (i) if, and which commodities are exchanged, and (ii) whether the exchange is affected by the availability of providers or commodities. This provides an exciting avenue for future research in wild birds.

## Conclusion

Our previous study on SC-cockatoos found that aggressive interactions are influenced by relative rank within the dyad, suggesting that knowledge about their place in the local hierarchy informs social decision within this species (Penndorf et al., 2025). We also previously found evidence that SC-cockatoos preferentially preen kin, as well as preening associates from other roosting groups when encountering each other in intermediate locations, suggesting that allopreening plays an important in maintaining bonds in this fission-fusion society (Penndorf et al., 2023). Here, we build on these findings with a fine-scale analysis that uncovers the importance of dominance rank in shaping the patterning of preening interactions. Our findings therefore add to the evidence for differentiated social relationships within SC-cockatoos, with individuals exhibiting a multi-faceted affiliative and aggressive response to other individuals that depends on roost membership, social network ties, rank and kinship. Our work also suggests intriguing directions for future work in this species that could add further valuable comparisons to primates and corvids.

## Authorship contributions

JP, LMA, and JMM conceived the study. JP, LF and LMA collated the social and morphometric data, and JP performed the analysis. All authors contributed to writing the manuscript.

## Acknowledgements

We would like to acknowledge the Gamaragal and Gadigal people as the traditional custodians of the land on which this study was conducted.

We thank Dr. Elizabeth Hobson for sharing the original code on which Figures 2b&c are based https://hobsonresearch.com/index.php/2015/04/10/attribute-ordered-networks/), Jana Horsch for assistance during the 2018 fieldwork, as well as Michael Chimento and Simeon Smeele for statistical advice.

## Funding

This study was partly funded through a Max Planck Society Group Leader Fellowship to LMA and by funding from the IMPRS for Organismal Biology to JP. JP and LMA are currently supported by the Swiss State Secretariat for Education, Research and Innovation (SERI) under contract number MB22.00056.

## Data availability

During the review process, all data and code are available on OSF (https://osf.io/zgwe8/overview?view_only=23295bfc9902474593ee3d6ca9a73548). Upon acceptance, this repository will be made public.

## Ethics statement

Research was approved by the Ethics Council of the Max Planck Society, Germany (application no. 2018_12). All procedures were approved by the ACEC (ACEC Project No. 19/2107), and were conducted under a NSW Scientific License (SL100107).

## Notes

### Competing Interest Statement

The authors have declared no competing interest.

### Summary of Updates

Main text of the manuscript has been revised.

